# Adaptive focused acoustics-integrated proteome profiling of macrophages uncovers low abundant proteins associated with immune homeostasis, inflammatory response, and transport

**DOI:** 10.64898/2026.05.26.728044

**Authors:** Jason A. McAlister, Michael Woods, Lia Abarzua, Sameer Vasantgadkar, Debadeep Bhattacharyya, Jennifer Geddes-McAlister

## Abstract

Efficient and reproducible protein extraction is a critical step in mass spectrometry-based proteomics workflows, particularly for complex host–pathogen systems where low-abundance immune-associated proteins are difficult to detect. Probe sonication methods used for cell lysis requiring mitigation of excessive heat generation, to prevent degradation of biologically important proteins, while also limiting throughput and potentially introducing sample-to-sample variability. In this study, we evaluated adaptive focused acoustics (AFA) technology as an alternative approach for macrophage lysis and protein extraction and digestion within a standard proteomics workflow coupled with mass spectrometry. We observed that AFA technology reduced hands-on processing times and overall workflow timelines and single-sample AFA technology improves proteome coverage, dynamic range, and reproducibility. We also evaluated multiplexed AFA technology for lysis, and we observed an exclusive macrophage proteome and influence on replicate reproducibility and dynamic range detection for low abundant proteins. Moreover, multiplexed AFA technology for macrophage lysis and digestion further increased protein identifications, replicate reproducibility, and dynamic range. Considering the AFA-exclusive proteome, 86 proteins were detected across all AFA-based lysis and digestion methods, including low-abundance proteins associated with macrophage homeostasis, inflammatory response, and transport. Together, these findings demonstrate that AFA technology enhances reproducibility, throughput, and proteome depth for macrophage protein extraction while enabling the detection of biologically relevant low-abundance immune-associated proteins. These improvements provide a strong foundation for future investigation of host–pathogen infection models, where pathogen-derived proteins remain challenging to detect within complex host proteomes.

## Introduction

The immune system relies on tightly regulated molecular networks to maintain cellular homeostasis, controlling inflammatory signaling, and mounting rapid responses to infection or environmental stress. These processes are governed by dynamic changes in pattern recognition receptors upon sensing a non-self molecular motif, such as a pathogen-associated molecular patterns (PAMPs), and synthesis and mobilization of communication signals. These activation processes and communication signals, beginning with innate immune system recognition manifest into adaptive immune responses for targeted and prolonged protection^1^. Critically, it is changes in protein abundance, localization, post-translational modification, and protein–protein interactions that shape immune cell activation and subsequent functions of the immune system^2,3^. While transcriptomic approaches have provided important insights into immune regulation, mRNA abundance does not always correlate with protein abundance or activity, particularly for rapidly regulated immune-associated pathways such as proteolysis, oxidative stress responses, cytokine signaling, and apoptosis^4–7^. As a result, direct measurement of proteins is important for understanding the functional state of immune cells and the molecular mechanisms underlying immune homeostasis and activation.

Mass spectrometry-based proteomics has emerged as a powerful systems-level approach for characterizing immune regulation through the unbiased detection and quantification of thousands of proteins within complex biological samples^2,8–10^. Advances in high-resolution instrumentation, peptide separation strategies, and computational analysis pipelines now enable deep profiling of immune-associated proteomes with improved sensitivity, dynamic range, and reproducibility. These technologies have facilitated the identification of signaling networks involved in innate and adaptive immunity, including activation and remodelling of pathways associated with immune cell activation, antigen presentation, cytokine production, metabolic reprogramming, oxidative stress, and programmed cell death^11–16^. Importantly, proteomics enables the detection of low-abundance regulatory proteins and dynamic molecular signatures that are often overlooked using conventional molecular approaches^17,18^. These molecular signatures have demonstrated roles in spatial and temporal tracking of the immune response during infection, and propose opportunities for improved diagnostics and prognostics, as well as exploration of therapeutic targets^19–22^.

Macrophages represent a central component of innate immunity and play critical roles in maintaining tissue homeostasis, coordinating inflammatory responses, and mediating host defense against invading pathogens^23–25^. Upon activation, macrophages undergo extensive proteome remodeling to support antimicrobial activity, immune signaling, phagocytosis, and cellular adaptation to stress^13,14,17^. These responses are highly dynamic and can vary substantially depending on environmental stimuli, infection state, host immune status, and cellular metabolism. Consequently, accurate characterization of macrophage proteomes requires robust sample preparation workflows capable of preserving protein integrity while maximizing reproducibility and proteome depth. For example, to tackle upstream sample preparation challenges of macrophages, we developed a microfluidics-enabled proteome profiling platform to distinguish remodeling of host and bacteria during infection^26^. This approach preserved morphological and viability states of macrophages and *Klebsiella pneumoniae* while revealing population-specific dynamics that modulate phagocytic evasion by the bacterium. However, conventional downstream protein extraction methods, including probe sonication, can introduce technical variability and excessive heat during cell lysis, potentially compromising the stability and detection of sensitive or low-abundance immune-associated proteins^27^.

Recent technological advances in sample preparation have focused on improving protein recovery, reducing variability, computational integration, automation, and increasing throughput for proteomics workflows^28–30^. Adaptive Focused Acoustics (AFA) technology provides a controlled acoustic energy-based approach for cell disruption and molecular processing that minimizes localized heating while enabling highly reproducible sample handling^31^. In addition to efficient cell lysis, AFA-based workflows may improve enzymatic digestion efficiency and support scalable multiplexed processing of biological samples. Despite these advantages, the application of AFA technology for investigating immune-associated proteome remodeling in macrophages remains underexplored. To address current limitations in macrophage proteomics workflows, we evaluated AFA as an alternative approach for protein extraction via cell lysis and enzymatic digestion within a standard mass spectrometry-based proteomics pipeline. Across both single-sample and multiplexed platforms, AFA processing reduced hands-on sample preparation time and improved overall workflow efficiency while maintaining robust proteome coverage. Notably, AFA-based workflows enhanced proteomic dynamic range and replicate reproducibility, supporting improved detection of low-abundance proteins. Multiplexed AFA lysis and digestion approaches further increased protein identifications and reproducibility across biological replicates while enabling scalable sample processing. Comparative analysis identified a subset of proteins uniquely detected using AFA-based workflows, including low-abundance proteins associated with macrophage homeostasis, inflammatory signaling, transport, and immune regulation. Together, these findings highlight the utility of AFA technology for improving proteome depth, reproducibility, and throughput in macrophage proteomics and establish a strong foundation for future studies investigating host–pathogen interactions.

## Material and Methods

### In vitro murine macrophage model

BALB/c immortalized murine macrophage cells (generously provided by Dr. Felix Meissner, University Hospital Bonn) were maintained in tissue culture plates using Dulbecco’s Modified Eagle’s Medium (DMEM) supplemented with 10% fetal bovine serum, 1% L-glutamine and 5% penicillin-streptomycin (pen-strep). Macrophages were seeded in 24-well plates at 5 × 10^4^ cells/well and incubated for 2 d at 37 °C with 5% CO_2_ to reach a confluence of 70-80% (approximately 2 × 10^5^ cells/mL). At confluence, macrophages were released from the plates with cold phosphate buffered saline (PBS) and cell scraping followed by centrifugation at 400 x g, washing twice with PBS, flash frozen in liquid nitrogen, and stored at -80 °C.

### Sample preparation for mass spectrometry analysis – cell lysis and protein extraction

Frozen macrophages were divided into 28 vials (approx. 1 x 10 ^6 cells/vial); four independent replicates were assigned to four processing platforms to assess cell lysis (16 samples total). For the probe sonication proteomics workflow, macrophages were prepared as previously described^32^. Briefly, macrophages were resuspended in Tris-HCl (pH 8.5) with protease inhibitor cocktail and sodium dodecyl sulphate [(SDS) 2% final] followed by probe sonication (30 s on/30 s off in ice bath, 5 cycles, 30% power; Thermo Fisher Scientific). For both single-sample AFA processing (E220) and multiplexed AFA processing (LE220Rsc and R230), macrophages were resuspended in Covaris Tissue Lysis Buffer (<2% SDS, no protease inhibitor cocktail; PN 520284) and were processed according to the enclosed conditions (Table 1). These samples were reduced with dithiothreitol (DTT; 10 mM final concentration) and alkylated with iodoacetamide (IAA; 55 mM final concentration) before acetone precipitation at −20 °C overnight. Precipitated proteins were collected by centrifugation, washed with 80% acetone twice, and solubilized in 8 M urea/40 mM HEPES. Protein concentration was quantified by tryptophan assay^33^, diluted in ammonium bicarbonate (ABC; 50 mM final concentration), normalized to 50 µg of protein, and digested with LysC-trypsin (Promega, protein:enzyme ratio, 50:1) overnight at room temperature. Digestion was stopped with 10% v/v trifluoroacetic acid and peptides were purified using C18 (three layers) Stop And Go Extraction tips^34^, vacuum dried, and stored at -20 °C until measurement.

**Table 1:**
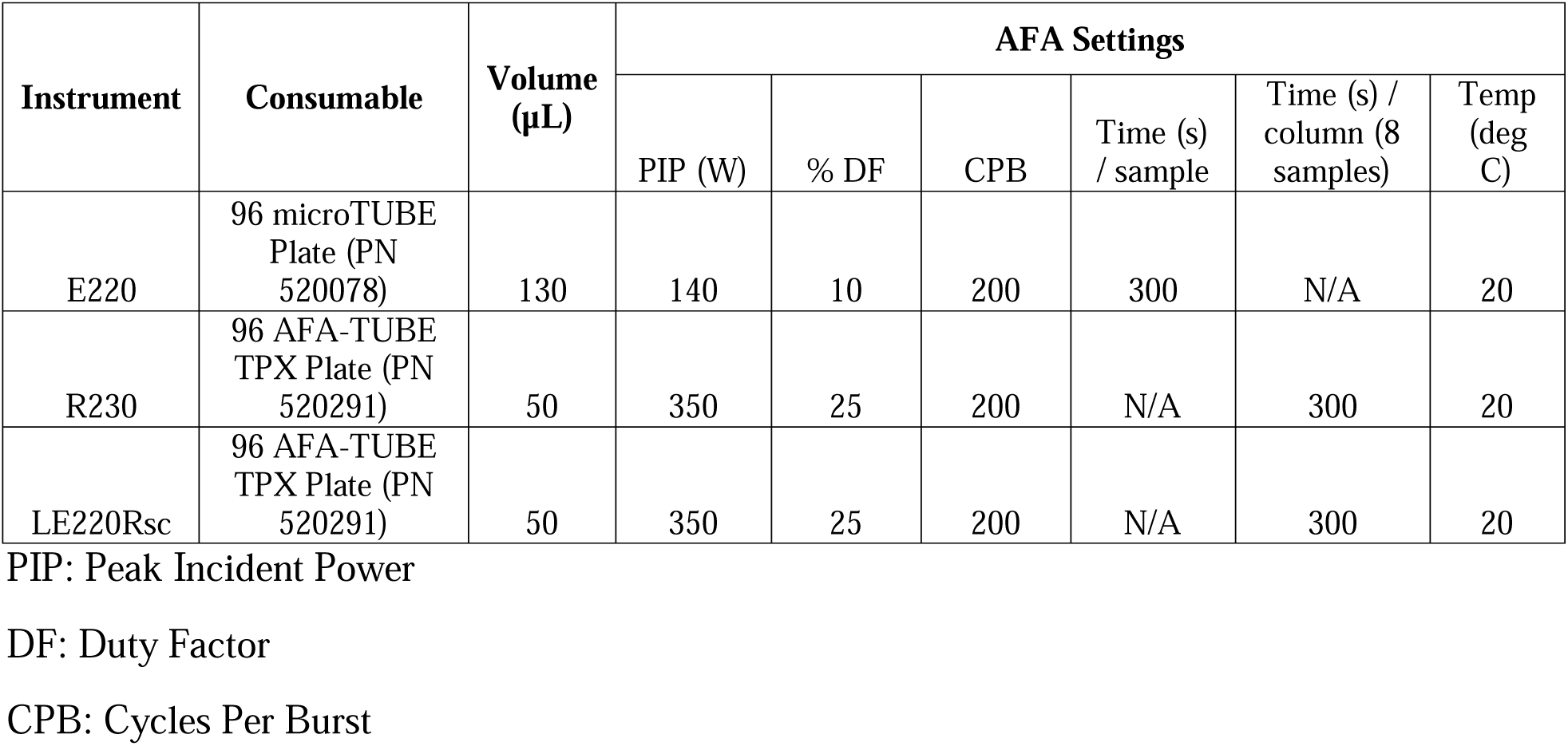
AFA settings used for macrophage sample processing.

### Sample preparation for mass spectrometry analysis – cell lysis and protein digestion

Of the remaining frozen macrophages (12 vials) four independent replicates were assigned to three processing platforms to assess cell lysis and digestion. Macrophages were prepared as described above with the following modifications. For the probe sonication proteomics workflow, samples were digested with LysC-trypsin (Promega, protein:enzyme ratio, 50:1) overnight at room temperature. For the AFA workflow, samples were lysed, reduced, and alkylated as described above using the LE220Rsc and R230 platforms (Table 1) followed by clean-up using the Protein Aggregation Capture and on-bead Trypsin Gold digestion (protein:enzyme ratio, 10:1) for 3 h or 1 h using the multiplex systems LE220Rsc and R230, respectively^35^. Peptides were purified using C18 (three layers) Stop And Go Extraction tips^34^, vacuum dried, and stored at -20 °C until measurement.

### Mass spectrometry

Purified peptides were resuspended in 0.1% formic acid and separated by a Vanquish neo ultra-high performance liquid chromatography system (Thermo Fisher Scientific, Bremen, Germany) using a 15 cm column with 75 μm inner diameter and 2 μm reverse-phase silica beads to separate peptides. Peptides were electrosprayed directly into the high-resolution Thermo Exploris 240 mass spectrometer (Thermo Fisher Scientific) using a linear gradient from 4% to 30% ACN in 0.1% formic acid over 120 min at a constant flow of 300 nl. The linear gradient was followed by a washout to 95% ACN to clean the column. The mass spectrometer was operated in a data-dependent mode, switching automatically between one full-scan and subsequent MS/MS scans of the most abundant peaks, with full-scans (400–1600 *m/z*) acquired.

### Mass spectrometry data processing

The unprocessed (.RAW) mass spectrometry files were analyzed using MaxQuant software (version 2.4.10.0)^36^. The derived peak list was searched with the incorporated Andromeda search engine against the reference *Mus musculus* (May 20, 2024; 54,510 sequences) from UniProt^37^. The following search parameters were set: trypsin enzyme specificity (arginine and lysine) with maximum 2 missed cleavages; minimum peptide length of seven amino acids; fixed modifications of carbamidomethylation of cysteine, variable modifications of methionine oxidation and N-acetylation of proteins. Peptide spectral matches were filtered using a target-decoy database at a false discovery rate (FDR) of 1% with a minimum of two peptides required for protein identification. Mass tolerance for precursor and fragment ions was set to 4.5 ppm and 20 ppm, respectively. Relative label-free quantification (LFQ) was enabled with MaxLFQ algorithm integrated into MaxQuant using a minimum ratio count of one^38^.

### Bioinformatics

Statistical analysis and visualization of the MaxQuant-processed data (proteingroups.txt file) were conducted using the Perseus software (version 1.6.2.2) and ProteoPlotter^39,40^. Hits to the reverse database, contaminants, and proteins solely identified by site were removed. LFQ intensities were log_2_ transformed and valid-value filtering (present in 3 of 4 replicates in at least one group) was performed. Missing values were imputed from a normal distribution (downshift of 1.8 standard deviations and a width of 0.3 standard deviations), followed by a Student’s *t*-test to determine proteins with significant differential abundance (*p*-value ≤ 0.05, S_0_ = 1) applying a 5% permutation-based FDR filter^41^. Resulting data was visualized with a Principal Component Analysis (PCA), hierarchical clustering (Pearson correlation) by Euclidean distance to showcase replicate reproducibility, in silico characterization of proteins of interest, including database searches (https://www.uniprot.org) and protein interactions depicted with STRING (https://string-db.org/). Graph Pad Prism (v. 10) was also used for data visualization.

## Results

### AFA technology reduced hands-on processing times and overall workflow timelines

AFA technology applies controlled acoustic energy to efficiently and reproducibly lyse biological samples, such as cells and tissues, for protein extraction^42,43^. Temperature control of the system aims to minimize localized sample heating, which can disrupt protein integrity, causing degradation of temperature sensitive biomolecules and reduced proteome dynamic range. Additionally, multiplexing capabilities reduce hands-on processing times, aiming to improve reproducibility and higher throughput compared to traditional (e.g., probe sonication) methods. To assess the impact of AFA technology within our established proteome profiling workflow, we prepared immortalized BALB/c macrophages and divided the cells across three workflows. First, we compared probe sonication of one sample at a time to a single-sample AFA workflow using the E220 (single-sample AFA) instrument. Under the test conditions, processing times of four biological samples is approximately 40 h regardless of the method with each step in the pipeline taking an equivalent amount of time (e.g., approx. 5 min for cell lysis/sample) (Fig. 1A). Next, we compared the probe sonication method with multiplexing AFA technologies (i.e., LE220Rsc and R230), which enabled processing of the macrophage samples in parallel, reducing the sample processing timeline by approx. 1 h for the specific workflow (Fig. 1B). Notably, LE220Rsc and R230 instruments process all samples at a time (up to 96 samples/h), reducing hands-on processing time and the overall workflow timeline.

**Figure 1.**
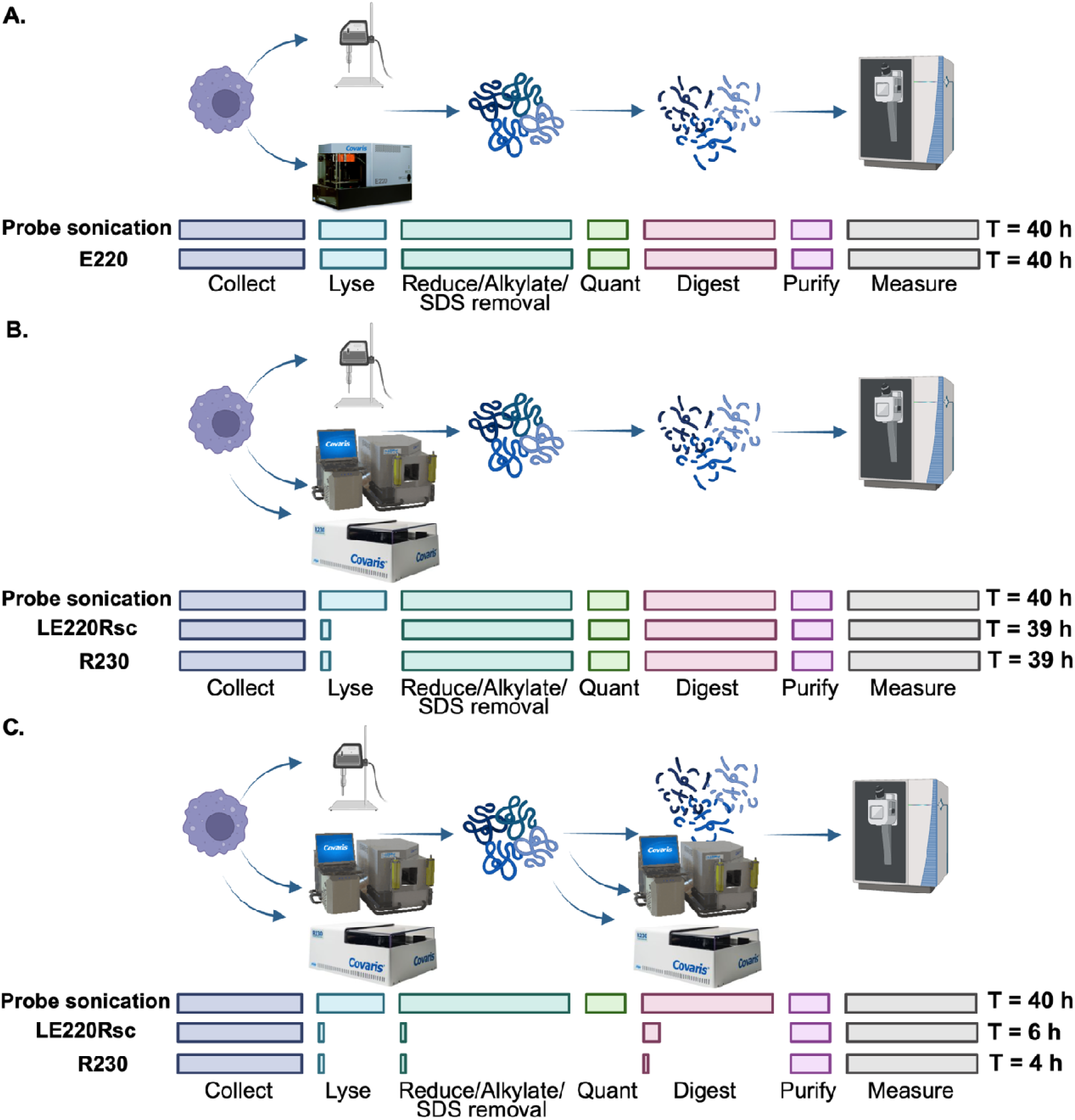
Sample preparation workflow and processing timelines. **A.** Probe sonication workflow with probe sonication compared to single-sample AFA technology (E220) followed by equivalent protein extraction and digestion, peptide purification, and mass spectrometry measurements. **B.** Probe sonication workflow with probe sonication compared to multiplexed AFA technology (LE220Rsc and R230) followed by equivalent protein extraction and digestion, peptide purification, and mass spectrometry measurements. **C.** Probe sonication workflow with probe sonication compared to multiplexed AFA technology (LE220Rsc and R230) followed by protein extraction, AFA-enabled multiplexed protein digestion, peptide purification, and mass spectrometry measurements. Workflow processing times presented at the end of each workflow. Experiments performed in four biological replicates.

Having performed cell lysis with probe sonication vs. AFA approaches while maintaining equivalent downstream sample processing workflows, we then evaluated a role for AFA technology on protein enzymatic digestion. Here, we applied the multiplexing AFA instruments (LE220Rsc and R230) for cell lysis, as described above, and for protein digestion, testing 3 and 1 h incubation times. We observed a substantial reduction in the overall workflow timeline from 40 h (probe sonication method with overnight enzymatic digestion) to 6 h using LE220Rsc with a 3 h digestion and 4 h using R230 with a 1 h digestion (Fig. 1C). Overall, application of AFA technology to lysis of macrophages and enzymatic digestion of proteins reduced hands-on sample processing times and workflow timelines by up to 60% over the probe sonication-based method.

### Single-sample AFA technology improves proteome coverage, dynamic range, and reproducibility

To assess the effect of cell lysis methods, such as probe sonication approach versus single-sample AFA technology (E220 instrument), we extracted and digested proteins from the samples and measured the proteomes on a mass spectrometer in data-dependent acquisition mode. Across four replicates for each condition, we identified an average of 3,238 (+/- 31) proteins using the probe sonication method compared to 3,299 (+/-16) proteins using the E220 AFA method (Fig. 2A). These data represent a significant increase (p-value = 0.0142) in protein identifications using the AFA method compared to the probe sonication approach. We visualized overlap and exclusive protein detection profiles between lysis methodologies, and we detected a common proteome of 2,976 proteins with an additional 266 (7%) proteins unique to the probe sonication method and 325 (9%) proteins unique to the E220 AFA method (Fig. 2B). A principal component analysis showed distinct clustering between the lysis methods (component 1, 49.13%) and separation by replicates (component 2, 11.23%) (Fig. 2C). As the distribution of replicates using the probe sonication method was broader than the AFA method, we performed a column correlation analysis by Euclidean distance of protein intensities, and we observed a significant increase (p-value <0.0008) in replicate reproducibility using the AFA technology (95% +/-0.5%) compared to the probe sonication approach (93% +/-0.8%) (Fig. 2D). A heatmap of hierarchical clustering by Euclidean distance confirmed clustering by lysis methodology and the enhanced replicate reproducibility using AFA technology (Fig. 2E). Next, we visualized dynamic range of proteins detected from each protocol and we observed a greater dynamic range, particularly for low abundant proteins following E220 AFA lysis (Fig. 2F). These observations were quantified with a significant increase (p-value = 0.0481) in mean normalized LFQ intensity and range of LFQ values for E220 lysis (0.03 +/-2.09) compared to the probe sonication method (-0.06 +/-2.07) (Fig. 2G). Taken together, these data demonstrate higher protein identifications, enhanced reproducibility among replicates, and increased dynamic range following macrophage lysis by single-sample AFA technology compared to probe sonication methods.

**Figure 2:**
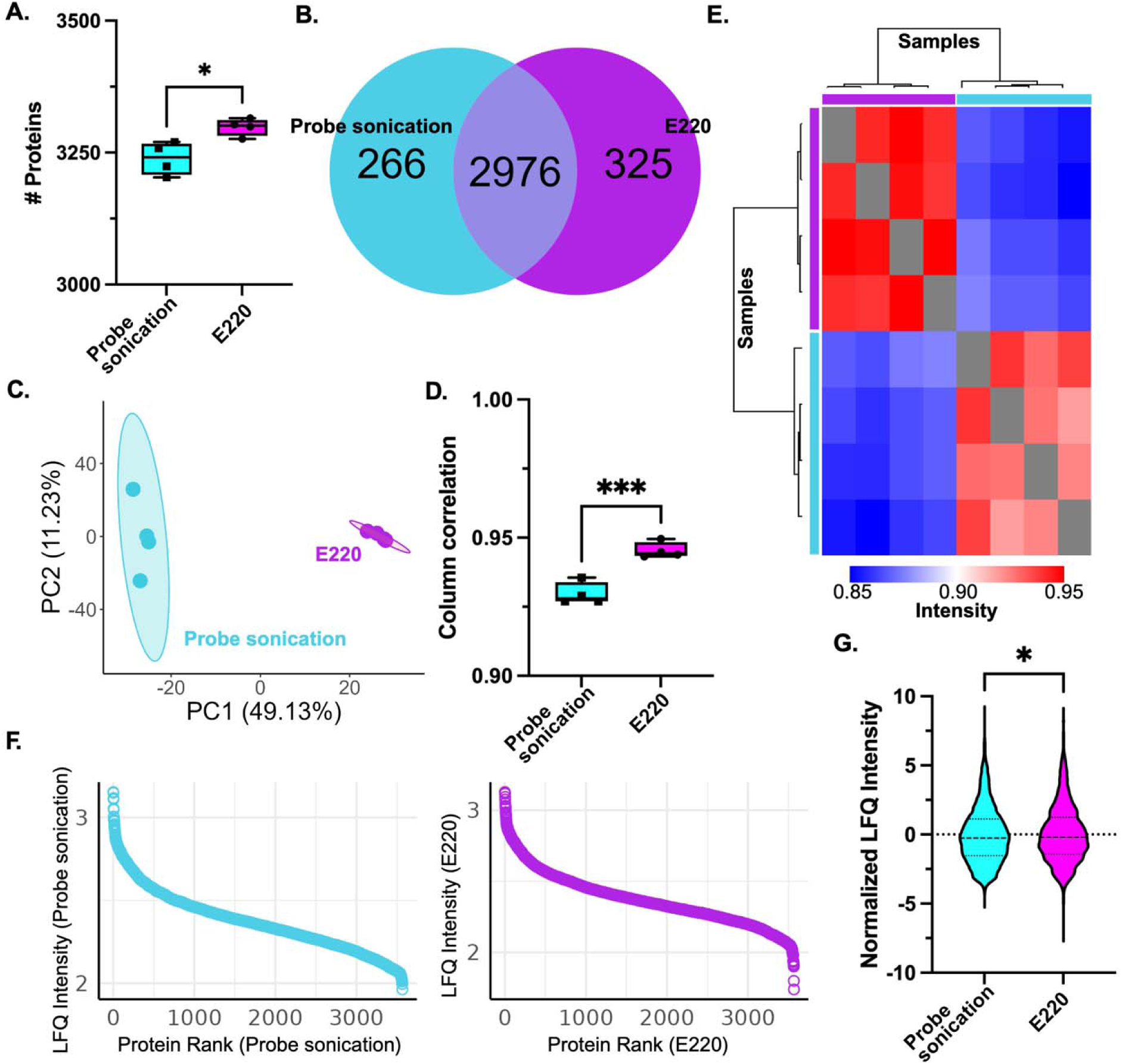
Proteome profiling of single-sample AFA compared to probe sonication macrophage lysis. **A.** Number of proteins detected between methods. **B.** Venn diagram for number of proteins common and unique to each method. **C.** Principal component analysis. **D.** Column correlation by Euclidean distance. **E.** Heatmap of hierarchical clustering by Euclidean distance for samples. E220 (purple) and Probe sonication (teal). **F.** S-curve of dynamic range across conditions. **G.** Normalized LFQ intensity median and value distribution. Statistical analysis by unpaired Student’s t-test *p-value < 0.05, ***p-value <0.005. Experiment performed in biological quadruplicate.

### Multiplexed AFA technology defines exclusive macrophage proteome and influences replicate reproducibility and dynamic range detection for low abundant proteins

Based on improvements in protein identifications, replicate reproducibility, and dynamic range of single-sample AFA lysis compared to the probe sonication workflow, we assessed the effect of multiplex sample AFA lysis. Here, we identified an average of 3,291 (+/-34) proteins using the probe sonication method compared to 3,358 (+/-28) proteins using the LE220Rsc (multiplexed), and 3,449 (+/-36) proteins using the R230 (multiplexed) AFA methods (Fig. 3A). Notably, one replicate of LE220Rsc was removed from the analysis due to low protein identifications and suggested technical error in sample processing. We observed a significant increase in proteins identifications for probe sonication vs. LE220Rsc (p-value = 0.028) and probe sonication vs. R230 (p-value = 0.0007). We visualized a common proteome of 2,607 proteins across all three lysis methods, with 119 (3%) proteins exclusively detected by probe sonication lysis compared to 693 (18%) proteins exclusively detected using the multiplexed AFA method (Fig. 3B). Notably, these numbers also represent AFA instrument-dependent proteomes (i.e., 119 proteins exclusive to LE220Rsc and 436 proteins exclusive to R230). A principal component analysis showed distinct clustering by lysis method with probe sonication and LE220Rsc processing separated from R230 (component 1, 58.27%) and separation by multiplexed vs. probe sonication (component 2, 13.03%) (Fig. 3C). To assess replicate reproducibility upon multiplexed AFA lysis, we performed a column correlation analysis by Euclidean distance of protein intensities. We observed a significant increase (p-value = 0.0009) in replicate reproducibility using the LE220Rsc AFA technology (95% +/-2.7%) compared to the probe sonication approach (93% +/-2.1%) and a significant decrease (p-value = 0.0256) using the R230 AFA technology (91% +/-1.7%) compared to the probe sonication approach (Fig. 3D). These reproducibility trends were further observed by a heatmap of hierarchical clustering by Euclidean distance (Fig. 3E). Next, we visualized the dynamic range of proteins detected from each protocol and we observed a greater dynamic range, particularly for low abundant proteins following multiplexed AFA (LE220Rsc and R230) lysis compared to the probe sonication method (Fig. 3F). These observations were quantified with a significant increase (p-value = 0.0458) in median normalized LFQ intensity and range of LFQ values for LE220Rsc lysis (0.03 +/- 1.93) compared to the probe sonication method (0.25 +/-1.91) but no significant difference (p-value = 0.4352) was observed for median normalized LFQ intensities by R230 lysis (0.18 +/-2.05) (Fig. 3G). Taken together, the multiplexed AFA lysis identified significantly more proteins (including exclusive identifications) compared to the probe sonication approach with replicate reproducibility and dynamic range differences dependent upon multiplexing approach.

**Figure 3:**
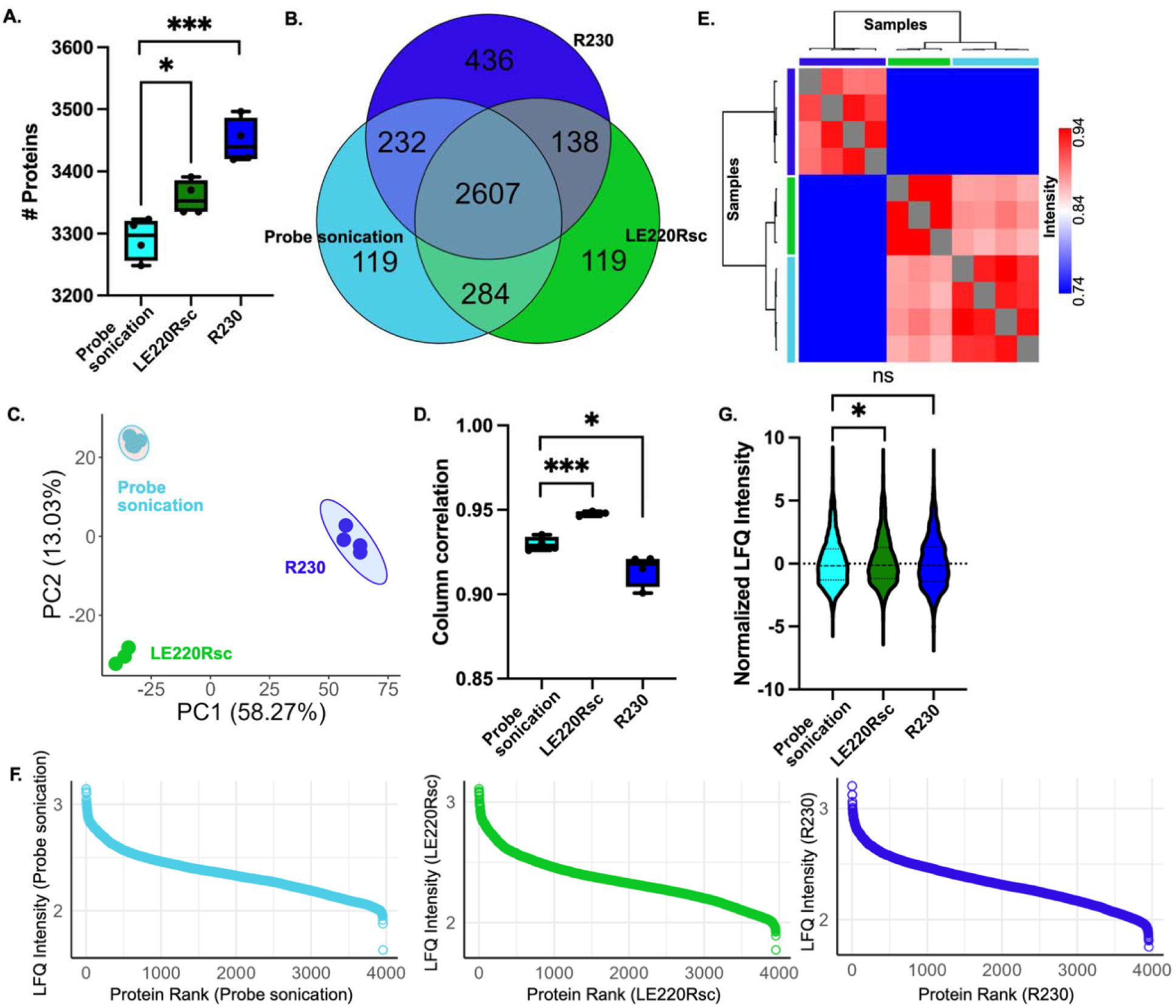
Proteome profiling of multiplexed AFA lysis compared to probe sonication macrophage lysis. **A.** Number of proteins detected among methods. **B.** Venn diagram for number of proteins common and unique to each method. **C.** Principal component analysis. **D.** Column correlation by Euclidean distance. **E.** Heatmap of hierarchical clustering by Euclidean distance for samples. R230 (dark blue), LE220Rsc (green), and probe sonication (teal). **F.** S- curve of dynamic range across conditions. **G.** Normalized LFQ intensity median and value distribution. Statistical analysis by unpaired Student’s t-test *p-value < 0.05, ***p-value <0.005. Experiment performed in biological quadruplicate.

### Multiplexed AFA technology for macrophage lysis and digestion continued to increase protein identifications, replicate reproducibility, and dynamic range

Given improvements in protein identifications using AFA technology compared to the probe sonication workflow, we aimed to evaluate if other steps in the proteome profiling pipeline could be optimized using AFA technology. Specifically, we focused on enzymatic digestion as the enzyme activity would likely benefit from controlled temperature and reduced digestion time would shorten the overall workflow timeline. By combining macrophage lysis and enzymatic protein digestion into the multiplexed AFA pipeline, we identified a significantly higher number of proteins for both LE220Rsc (p-value = 0.0001; 3,400 +/- 10) and R230 (p-value = 0.0001; 3,406 +/- 17) instruments compared to the probe sonication (3,253 +/-28) approach (Fig. 4A). We defined a common proteome of 2,918 across the three methods with 220 (5.9%) proteins exclusive to the probe sonication approach and 492 (13.1%) exclusive to the AFA platforms (Fig. 4B). Notably, AFA instrument-dependent proteomes were observed (i.e., 67 proteins exclusive to LE220Rsc and 83 proteins exclusive to R230). A principal component analysis showed distinct clustering by method with probe sonication lysis and digestion separating by multiplexed AFA lysis and digestion (component 1, 49.23%) and separation by replicates (component 2, 7.61%) (Fig. 4C). Next, we evaluated replicate reproducibility amongst the samples, and we observed the highest values across experiments performed within this study. Specifically, we observed a significant increase in reproducibility upon use of the multiplexed LE220Rsc AFA system (p-value = 0.0147; 95% +/- 0.7%) and R230 AFA system (p-value = 0.0052; 94.2% +/- 0.8%) compared to the probe sonication method (93.1% +/- 0.8%) (Fig. 4D). These reproducibility trends were observed by a heatmap of hierarchical clustering by Euclidean distance (Fig. 4E). We also visualized the dynamic range distribution of proteins detected from each protocol and, consistent with the previous analysis, we observed a greater dynamic range, particularly for low abundant proteins following multiplexed AFA (LE220Rsc and R230) lysis and digestion compared to the probe sonication method (Fig. 4F). Quantification of these observations showed a significant increase (p-value < 0.0009) in median normalized LFQ intensity and range of LFQ values for LE220Rsc (-0.03 +/- 2.10) and R230 (-0.04 +/-2.08) lysis and digestion compared to the probe sonication method (-0.19 +/-2.13) (Fig. 4G). Taken together, the multiplexed AFA lysis and digestion protocol continued to increase protein identifications, replicate reproducibility, and dynamic range compared to the probe sonication method.

**Figure 4:**
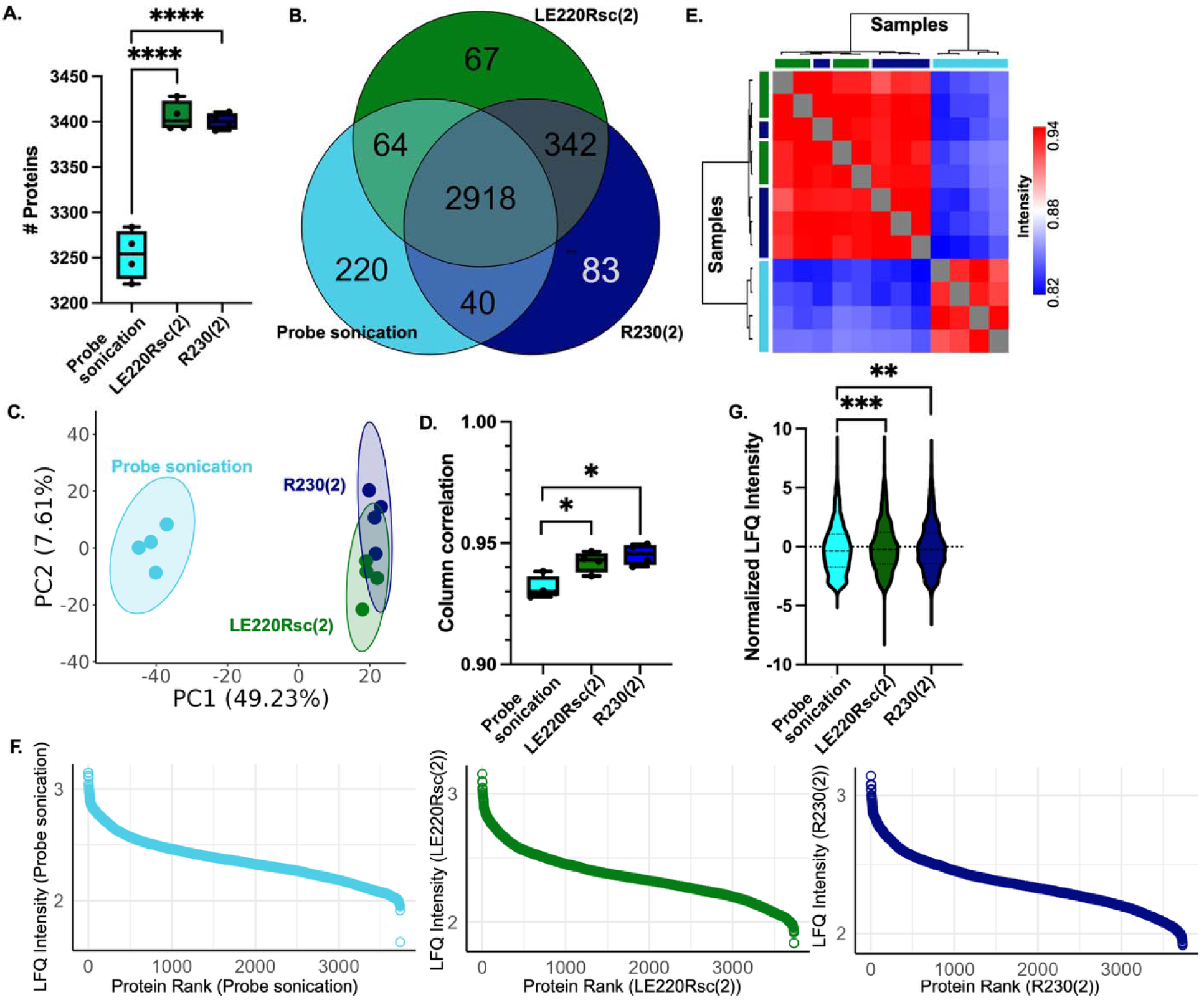
Proteome profiling of multiplexed AFA lysis and digestion compared to probe sonication workflow. **A.** Number of proteins detected among methods. **B.** Venn diagram for number of proteins common and unique to each method. **C.** Principal component analysis. **D.** Column correlation by Euclidean distance. **E.** Heatmap of hierarchical clustering by Euclidean distance for samples. R230(2) (dark blue), LE220Rsc(2) (dark green), and probe sonication (teal). **F.** S-curve of dynamic range across conditions. **G.** Normalized LFQ intensity median and value distribution. Statistical analysis by unpaired Student’s t-test *p-value < 0.05, **p-value < 0.01, ***p-value <0.005. Experiment performed in biological quadruplicate. (2) depicts that both cell lysis and enzyme digestion were performed with the indicated AFA instruments.

### AFA-exclusive proteins show enrichment across macrophage homeostasis, inflammatory response, and transport

Based on the observed increased protein identifications using AFA technology for macrophage lysis and digestion, we aimed to explore the functional roles of proteins exclusive to these profiles. Across all six experimental conditions, we observed 2,483 proteins commonly detected, and 1,022 proteins that were exclusive all combinations of AFA-treated samples (Fig. 5A). Of these, given technical and instrument differences across AFA platforms, we focused on 86 proteins commonly detected across all AFA experiments but absent from the probe sonication approach. Given our observation of increased protein dynamic range using AFA technology, we hypothesized that the median LFQ intensities of proteins exclusive to AFA treatment would be lower than that of all proteins detected. Our hypothesis was supported with a significant reduction (p-value < 0.0001) in mean log_2_ intensity for the 86 prioritized proteins (24.83 +/-1.00) compared to all other proteins (26.12 +/- 0.39) (Fig. 5B). Next, we investigated protein-protein interactions and defined networks among the 86 proteins using STRING database^44^. By MCL (Markov Cluster Algorithm) clustering, we observed 17 protein clusters with those having defined functional descriptions labeled (Fig. 5C; Fig. S1A). Notably, we observed clustering of proteins associated with cytokinesis, protein degradation (aggrephagy, Zn-finger N-recognin), transport, inflammatory, signaling, and translational regulation (large ribosomal subunit, endoribonuclease, origin replication complex). An enrichment plot based on Gene Ontology Biological Processes highlighted vesicle mediated transport to the plasma membrane, secretion by cell, and establishment of organelle localization as the top three categories (Fig. 5D). Additionally, enrichment by Gene Ontology Cellular Component defined cytosol, intracellular anatomical structure, and intracellular organelle among the top three categories (Fig. S1B). We also observed 48 proteins that were exclusive to the probe sonciation proteomics method (i.e., not identified across the AFA-processed samples) (Fig. S2A). Of these proteins, three clusters were identified by MCL clustering with one cluster functionally defined with as peroxisome biogenesis; no enrichment by Gene Ontology categories was observed (Fig. S2B). Taken together, these findings support an exclusive proteome detected with the application of single-sample and multiplexed AFA technology for macrophage lysis and digestions. Biological functions of these AFA-exclusive proteins span an array of processes but emphasize enrichment of proteins associated with macrophage homeostasis, inflammatory response, and transport.

**Figure 5:**
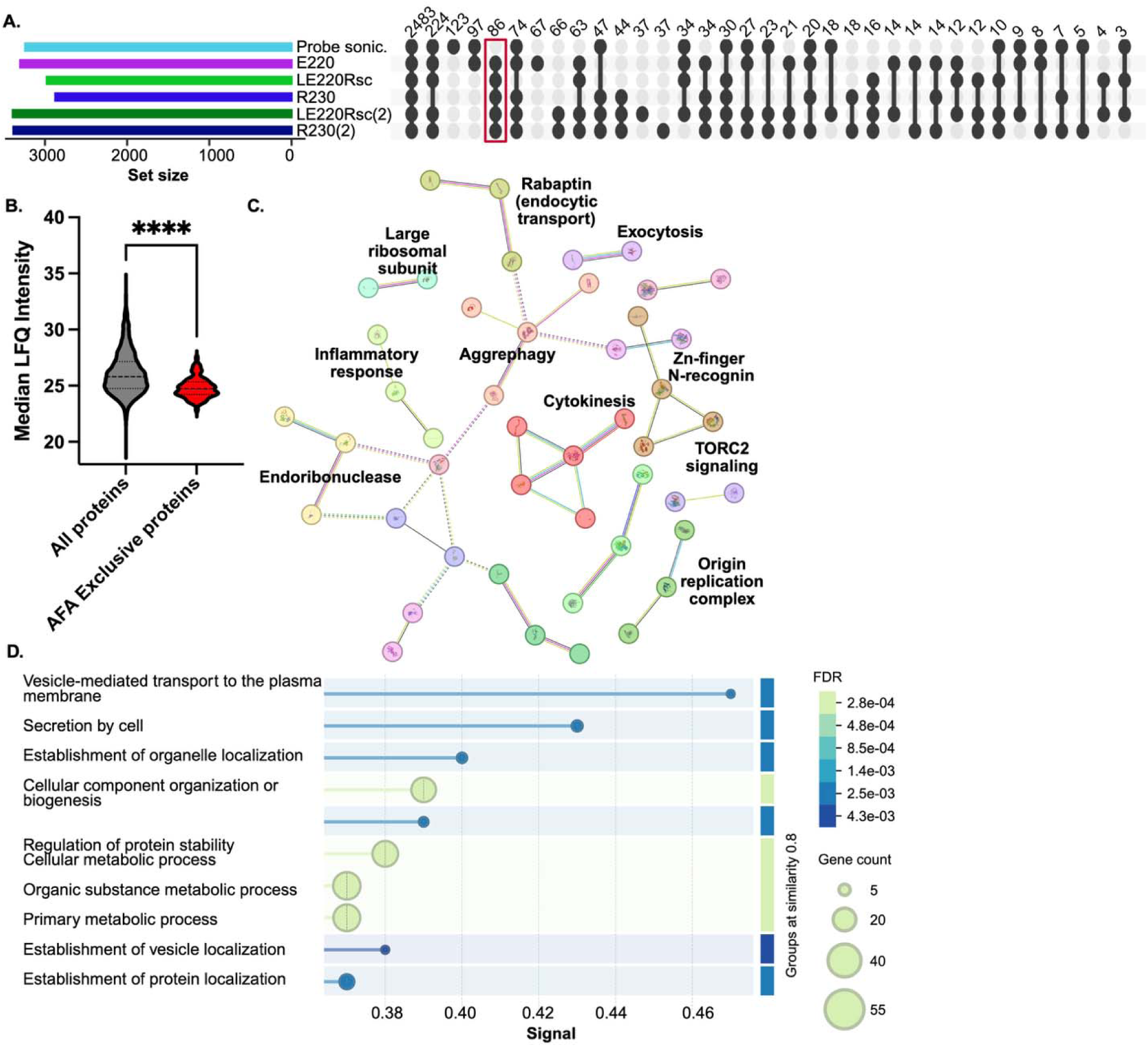
Functional assessment of an AFA-exclusive proteome. **A.** UpSet plot for number of proteins detected across the six experimental conditions. Red box highlights proteins (86) exclusively detected using AFA technology for macrophage lysis and digestion. (2) depicts that both cell lysis and enzyme digestion were performed with the indicated AFA instruments. **B.** Median LFQ intensity of AFA-exclusive proteins (86) and all other detected proteins (4,098). Statistical analysis by unpaired Student’s t-test ****p-value < 0.0001. **C.** Protein-protein interaction network by STRING database. By MCL clustering, 17 clusters were defined – clusters with functional annotation labeled; non-interacting proteins removed from network visual. MCL clustering by inflation parameter = 3, medium confidence interaction score = 0.400. **D.** Enrichment map by Gene Ontology Biological Processes. Terms grouped by >0.8 similarity, terms sorted by signal (>0.25) and 5% FDR. Experiment performed in biological quadruplicate.

## Discussion

This study evaluated AFA technology as an alternative strategy for macrophage lysis and enzymatic digestion within a standard mass spectrometry-based proteomics workflow. Building on the need for reproducible and scalable approaches to investigate immune homeostasis, regulation, and activation, we demonstrate that AFA-based processing improves workflow efficiency while maintaining robust proteome coverage and strong quantitative reproducibility. Across both single-sample and multiplexed instrumentation platforms, AFA technology enhanced protein identifications, replicate reproducibility, and proteomic dynamic range, while reducing hands-on processing time. Importantly, AFA workflows enabled the detection of low-abundance proteins associated with macrophage homeostasis, inflammatory response, transport, and signaling that were not detected using probe sonication workflows. More broadly, improvements in reproducibility, scalability, and proteome depth are increasingly important as proteomics workflows move toward high-throughput and translational applications in immunology, infectious disease, and precision medicine.

Although the E220 single-sample AFA platform resulted in a statistically significant increase in protein identifications and replicate reproducibility compared to the probe sonication workflow, both methods demonstrated strong overall performance, with more than 3,000 proteins identified and replicate reproducibility values exceeding 90%. Similarly, multiplexed AFA approaches and probe sonication workflows shared a large common proteome and exhibited high reproducibility across replicates. These findings indicate that probe sonication workflows remain effective for broad macrophage proteome profiling, while AFA technology provides incremental, yet meaningful, improvements in consistency and proteome depth. Importantly, multiplexed AFA processing produced an approximately 18% increase in unique protein identifications relative to the probe sonication workflow, highlighting a substantial enhancement in proteome coverage when scaling sample throughput. The observed differences between multiplexed instrumentation platforms suggest that acoustic energy delivery influences proteomic outcomes. Notably, differences in the number of proteins identified across comparisons with probe sonication proteomes likely reflect stringent valid value filtering criteria requiring protein detection in three out of four replicates within at least one experimental group.

The continued improvement in protein identification rates, replicate reproducibility, and dynamic range observed following integration of AFA-assisted digestion further supports the utility of acoustic processing across multiple stages of the proteomics workflow. In addition to reducing overall workflow timelines, AFA-assisted digestion may enhance enzymatic accessibility and improve sample homogenization, thereby contributing to reduced technical variability among replicates. Similar observations have been reported in studies evaluating controlled acoustic energy for biomolecule extraction and sample processing, where temperature-controlled sonication reduced sample degradation and improved analytical consistency relative to probe-based methods^31,42,43^. The reduced localized heating associated with AFA technology may be particularly important for preservation of thermally sensitive and low-abundance proteins mediating signaling, degradative processes, and exocytosis and endocytosis, which are rapidly turned over or highly dynamic during macrophage activation^45–48^.

Proteins exclusively identified using AFA-based lysis and digestion approaches revealed enrichment of biological pathways associated with cytokinesis, proteostasis, vesicle trafficking, inflammatory signaling, transport, and translational regulation. Within the protein–protein interaction networks, enrichment of proteins associated with aggrephagy, zinc-finger N-recognin-mediated protein degradation, ribosomal regulation, endoribonuclease activity, and origin recognition complex biology suggests improved detection of tightly regulated cellular maintenance and stress-response pathways^49^. These processes are central to macrophage homeostasis and activation, where rapid remodeling of protein turnover, translation, and intracellular trafficking supports immune adaptation to environmental stimuli and infection. Similarly, enrichment of Gene Ontology Biological Processes associated with vesicle-mediated transport to the plasma membrane, secretion, and organelle localization likely reflects improved extraction or preservation of membrane-associated and trafficking-associated proteins that are often challenging to detect using probe sonication workflows. Macrophages rely heavily on vesicular transport pathways for cytokine secretion, phagosome maturation, antigen presentation, and intracellular signaling, processes that involve dynamic membrane remodeling and low-abundance regulatory complexes^50–52^. Because probe sonication can introduce excessive localized heating and mechanical variability, these proteins may be more susceptible to degradation or inconsistent extraction. In contrast, the controlled and non-contact acoustic energy generated during AFA processing may better preserve protein integrity and support more uniform disruption of intracellular and membrane-associated compartments. Enhanced recovery of proteins involved in proteostasis and vesicle trafficking may therefore provide deeper biological insight into macrophage immune regulation and inflammatory signaling states. Alternative technical considerations include the use of a Tris-HCl buffer with 2% SDS and protease inhibitor cocktail for the probe sonication lysis buffer, whereas when testing the AFA instruments for cell lysis, the Covaris Tissue Lysis Buffer (<2% SDS, no protease inhibitor cocktail) was used, which may influence solubility of membrane- or lipid-associated proteins and protease activity. To further assess the potential role of lysis buffers and digestion enzymes, future analyses can tease apart common and exclusive proteins across the AFA platforms tested; however, notably, the use of lower concentration of SDS, lack of a protease inhibitor cocktail, and trypsin lacking Lys-C for AFA-processed samples still resulted in higher protein identifications compared to the probe sonication approach. Additional technical considerations include the use protein aggregation capture and on-bead digestion performed using LE220Rsc and R230 platforms, which substantially reduce the overall workflow timelines but also introduce new technical variables. Future experiments will incorporate the protein aggregation capture and on-bead digest within the probe sonication workflow to reduce overall processing times and assess direct improvements using these alternative strategies within the probe sonication workflow.

Several opportunities remain to further optimize and accelerate the workflow described in this study. The current probe sonication workflow includes detergent-based lysis and SDS removal through acetone precipitation, both of which substantially contribute to processing time and sample handling complexity. Future studies could evaluate detergent-free extraction approaches or alternative cleanup strategies, such as S-Trap-based processing, to simplify sample preparation and reduce losses associated with precipitation workflows^53,54^. Digestion timelines may also be substantially reduced, as recent studies have demonstrated efficient proteolysis within approximately one hour while maintaining proteome depth and quantitative reproducibility^55^. Additional improvements may be achieved through advances in mass spectrometry instrumentation and acquisition strategies. Emerging platforms, such as the Orbitrap Astral Zoom provide substantially increased scan speed, ion utilization efficiency, and proteome depth, particularly when coupled with data-independent acquisition workflows^56^. Integration of these approaches could further reduce instrument acquisition time while increasing sensitivity for low-abundance proteins^21,57^. Moreover, because probe sonication may represent an excessively harsh lysis strategy for macrophage proteomics, future comparisons with more comparable approaches, such as waterbath sonication, may better define the specific advantages of AFA processing independent of heat-associated degradation effects. Direct comparisons of digestion time points between probe sonication and AFA-assisted workflows would also strengthen future benchmarking studies. Finally, extending these workflows into complex infection models involving macrophages and pathogens will provide important opportunities to investigate immune activation, inflammatory signaling, vesicle trafficking, and pathogen-associated proteome remodeling within biologically relevant host–pathogen systems.

## Author Contributions

J.A.M., L.A., S.V., D.B., & J.G.-M. conceptualized and designed the study. J.A.M., M.W., L.A., & S.V. performed experiments. J.A.M. prepared and processed samples for mass spectrometry. J.A.M. & J.G.-M. performed data analysis and designed and developed figures. J.G.-M. wrote and edited the manuscript. All authors have read and approved the submitted manuscript.

## Acknowledgments

Thank you to members of the Geddes-McAlister lab for their informative and constructive feedback on project design and manuscript preparation. The authors thank Dr. Dyanne Brewer of the Advanced Analysis Centre – Mass Spectrometry Facility at the University of Guelph for operation of the mass spectrometer, and Narendra Singh Yadav and Jeffrey Seitz of D-Mark Biosciences for their collaboration.

## Data Availability

The. RAW and affiliated files will be deposited into the publicly available PRIDE partner database for the ProteomeXchange consortium.

