## Supplementary material for "Adaptive focused acoustics-integrated proteome profiling of macrophages uncovers low abundant proteins associated with immune homeostasis, inflammatory response, and transport": Supp. Fig.

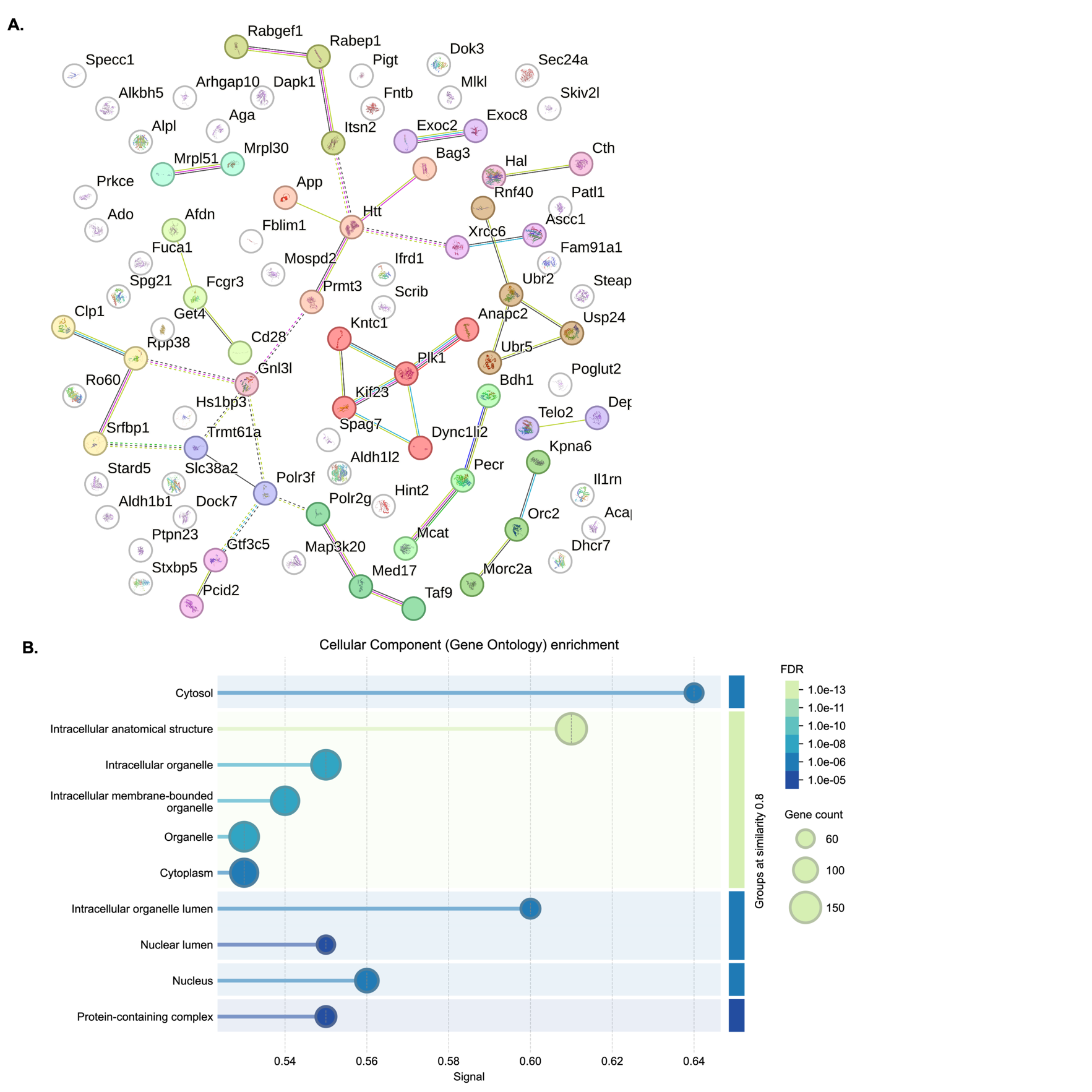

**Figure S1: Expanded functional assessment of an AFA-exclusive proteome. A.** Protein-protein interaction network by STRING database. By MCL clustering, 17 clusters were defined – all proteins maintained within the network. MCL clustering by inflation parameter = 3, medium confidence interaction score = 0.400. **B.** Enrichment map by Gene Ontology Cellular Compartment. Terms grouped by >0.8 similarity, terms sorted by signal (>0.25) and 5% FDR. Experiment performed in biological quadruplicate.

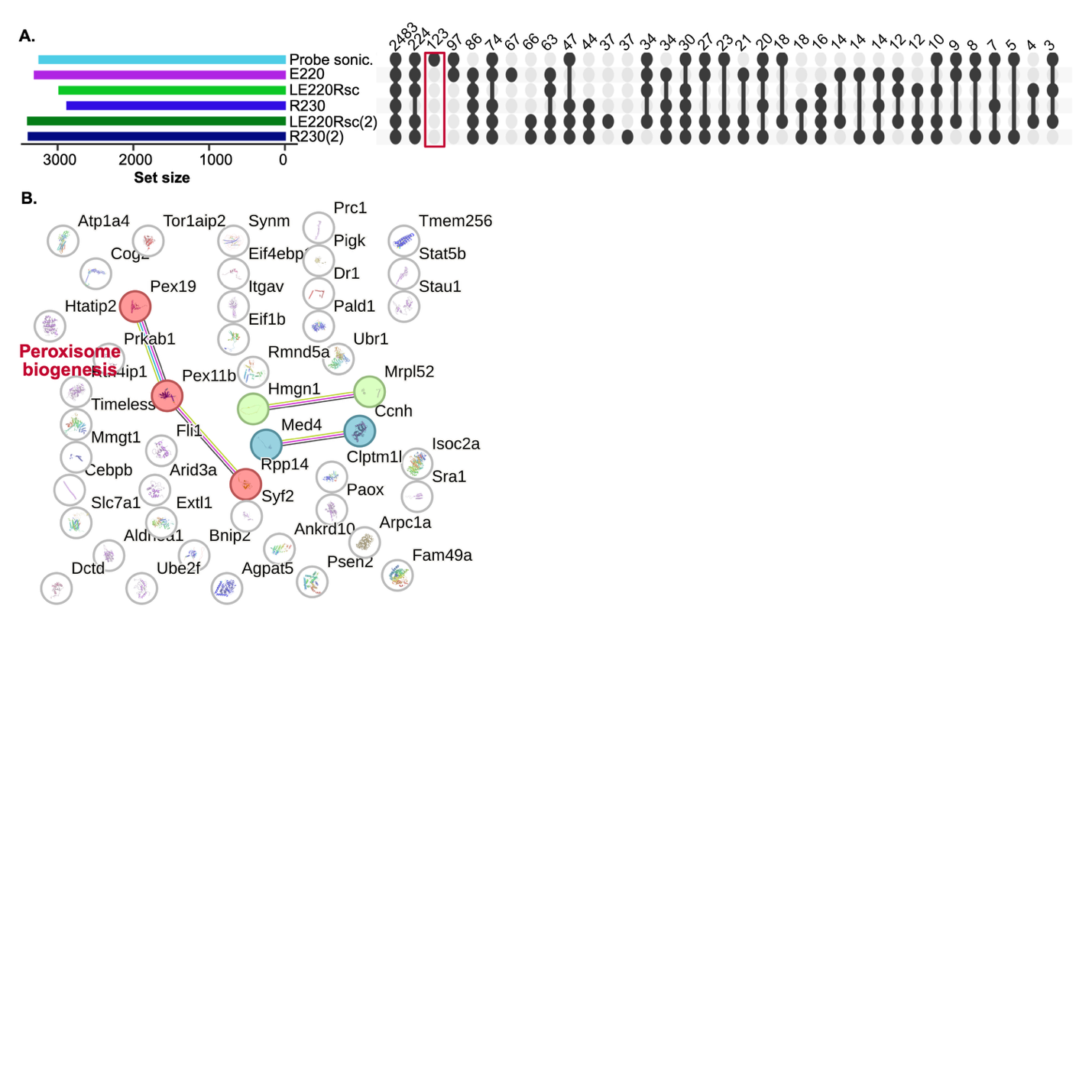

**Figure S2: Expanded functional assessment of a probe-sonication-exclusive proteome. A.** UpSet plot for number of proteins detected across the six experimental conditions. Red box highlights proteins (48) exclusively detected in probe sonication samples. **B.** Protein-protein interaction network by STRING database. By MCL clustering, 3 clusters were defined – all proteins maintained within the network. MCL clustering by inflation parameter = 3, medium confidence interaction score = 0.400. Experiment performed in biological quadruplicate.
